# Energy saving in vision at the first synapse: The ON and OFF pathways measure temporal differences

**DOI:** 10.1101/225557

**Authors:** Bart M. ter Haar Romeny

## Abstract

The inner plexiform layer (IPL) of mammalian retina has a precise bisublaminar organization in an inner on- and an outer off-layer, innervated by spatially segregated on- and off-cone bipolar cell inputs. Also, the processes of starburst amacrine cells are segregated into on and off sublaminae of the IPL. Distances between overlapping on-off pair retinal ganglion cell dendritic tree centers are markedly smaller than between on-on or off-off centers, indicating simultaneously sampling the same space. Despite dekades of research, no good model exists for the role of the on- and off pathways. Here I propose that the on- and off pairs are temporally subtracted, with one channel delayed in time, likely in a higher cortical center. The on- and off receptive fields give at every retinal location an I+ and I-signal, where I is intensity, velocity, color. Subsequent frame subtraction is a basis function of every surveillance camera for vision, and in MPEG video/sound compression. The model explains the many phenomena observed when the retinal image is stabilized. The separation of layers in the LGN fits with the notion of a time delay at higher cortical level. The directionalty observed in micro-saccades is typically perpendicular to the main edges in the scene. Precise measurement of spatio-temporal receptive field kernels shows that time is processed in the visual system as a real-time process, i.e. with a logarithmic time axis. As only contours and textures are transmitted, it is a very effective design strategy of the visual system to conserve energy, in a brain that typically uses 25 Watt and very low neuron firing frequencies. The higher visual centers perform the fill-in (inpainting) with such efficiency, that the subtraction always goes unnoticed.

## 1 Introduction

In all vertebrate species and insects on- and off-bipolar cell inputs are segregated in the inner plexiform layer (IPL) of the retina: synapses of on bipolar cells connect in the proximal half and those of the off bipolar cells connect in the distal half of the IPL [17,36,43,16]. The segregation of on- and off-pathways is one of the oldest findings in retinal physiology [29, 52], and is described in virtually every textbook [40, 27, 33]. It is extremely basic: both in vertebrate and fly photoreceptors the signal is split into on- and off-channels at the first synapse.

Many aspects have been thoroughly investigated, and its anatomical connections from stellate amacrine cells (SAC), bipolar and ganglion cells has been mapped from electron microscopy [26] to connectomic connectivity at the level of individual connections [22, 24], but the specific functional role of the two pathways has been unclear [7,45,46,33]. Westheimer suggested that the birather than a unidirectional nature of the retinal output has advantages in allowing small signals to remain prominent over a greater dynamic range [53]. Recently a role in HassensteinReichardt style motion detection was proposed for the on-off pathways in the fruitfly [23]. But the most widely accepted view is the simultaneous sensitivity for both dimming and brightening [27, 40].

Around 30 different functional and anatomical types of retinal ganglion cells (RGC) have been identified [3,33,42,12]. Profound dendritic redistributions in the IPL into the on- and off-lamina are seen after eye opening [54], while preventing the stratification with 2-amino-4-phosphonobutyric acid (APB, [31]) could be reversed by normal visual experience [7], indicating that visual stimulation is needed for the formation of the on-and off sublamina.

The number of on- and off RGCs is equally distributed: 50% is on, 50% is off. The mosaics of on-center and off-center retinal ganglion cells (RGCs) exhibit a complete tiling of the full retina [8]. In a careful mapping of the morphology and topography of the on- and off retinal alpha cells in the cat retina, Wässle found the distance between on-center and off-center ganglion cell bodies significantly smaller that the average distance between on-center and off-center separately [51]. Functionally, this indicates that the on- and off-mosaics need to sample the same spatially synchronized visual input. There are two midget cells, an on- and an off-cell, for every cone [40].

The on-center and the off-center RF give a representation of +*I* and –*I* pathways, with I the intensity, color or velocity. The pathways are subtracted, where one pathways is slightly delayed relative to the other. The delay may happen at cortical level. This gives the cortical circuits control over the delay, to make it adaptive to the task or the history of the signal. The measurement of 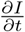 is likely to happen at a range of temporal scales.

This new realization of the function of the on-off pathways explains the effects seen during stabilized retinal images, of the 50%/50% on-off RF distribution, the segregation of layers in the LGN, the purpose of the direction of micro- and normal saccades, and is a very effective way to save energy. Subtraction of subsequent frames is the most often used image processing mode in surveillance cameras. When there is no motion, the signal is an energy-effective zero signal, and motion automatically highlights only the edge locations where the motion occurs.

This function of the on- and off pathways has not been described before. As we make eye- and head motions continously, and there is typically motion in the observed scene, we are completely not aware that we use a subsequent frame subtraction camera continuously. The brain is capable of a flawless and real-time filling-in process.

The paper is organized as follows. In the next section we justify the proposed model from the perspective of computer vision experience, energy efficiency and anatomical evidence. The role of on- and off channels measuring the temporal derivative with a delay at cortical level fits with the separation of pathways in the layers of the LGN, which is discussed in section 3. In section 4 we discuss the historic discoveries from stabilized retinal images. The purpose of saccades direction and the renewed interest in micro-saccades and attentional effects is described in section 5. In section 6 we discuss the physical necessity of a logarithmic time scale of real-time temporal measurements. The very robust higher level fill-in or inpainting process gives rise to an eloquent internal representation of the 3D visual world. In section 7 we make the comparison of a foveated scanning eye with the lidar-scanning robotics technique of V-SLAM (Visual Simultaneous Localization and Mapping). The paper is concluded with a conclusion and discussion.

## 2 Subsequent frame subtraction

The on- (+*I*) and off- (–*I*) channels carry positive and negative spatial signal fields for intensity/color (midget RGCs), and velocity (parasol RGCs). Adding the channels after a suitable temporal delay of one of the channels gives the temporal derivative 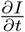, likely at the level of the cortex. The measurement of the first temporal derivative by cortical receptive fields was first shown by DeAngelis et al., applying a reverse correlation technique to map the 2D post-stimulus time histograms at a range of delays after the stimulus presented at random positions in the RF area on the retina [11].

A subtraction of subsequent frames (‘frame differencing’) is a common function in video surveillance cameras, and leads to highlighting of the areas that change intensity under motion of objects. See fig. 1. It divides an image frame into changed and unchanged regions. The changed region is associated only with the moving object. This Mathematica 11 command shows the effect in real-time:

**Fig. 1.**
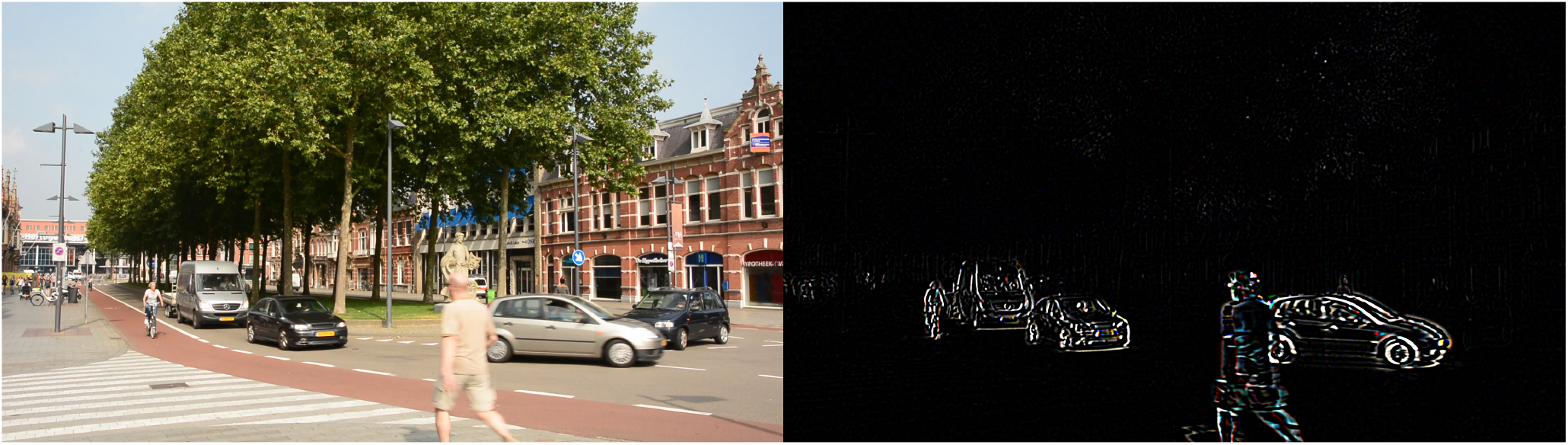
On-off pathways perform delayed subtraction. Compare this with subsequent frame differencing in real-time frame-differencing surveillance video cameras, highlighting only the moving contours.

Dynamic[ImageDifference@@CurrentImage[2]].

## 3 Layers of the LGN - Arguments for cortical time delay

The segregation into on- and off-pathways in the visual pathway is at the earliest stage possible: each foveal cone projects to two midget ganglion cells, an on-off pair, by their on- resp. off-bipolar cells [40].

Axons of all retinal midget ganglion cells project to only the LGN, where they end in the segregated parvo-cellular layers. The on-center midget cells terminate in layer *P*_4_ and *P*_3_, the off-center midget cells terminate in layers *P*_6_ and *P*_5_ [40]. In the ferret it was found that on-center and off-center axons end in different sublayers of layer A in the LGN [41]. In contrast, the on-off retinal ganglion cells project extensively to the superior colliculus [47]. The first parvo pair may transfer the pathway of the midget cells to the geometric analysis kernels in V1, to process texture, color, motion and disparity, while the second parvo pair may transmit the time successive difference. The magno pair transmits the pathways of the motion sensitive ganglion cells. This continued segregation of the pathways through the LGN indicates that the delay of one channel (intuitively, and from evolutionary perspective most likely the off-channel) and subsequent addition of the +*I* and –*I*–*δt* channels is not effectuated in the retina or LGN, but in the visual cortex. It is possible that a range of delays is incorporated, under adaptive control in the cortex, to perceptually group the incoming multi-scale spatio-temporal differential structure.

The general strategy of the nervous system, and in particular vision, to only deal with variations is a highly effective mechanism to save energy by reducing the number of active cells and active transfer lines. It needs a sophisticated internal representation in memory, which is continuously being updated (see section7).

In some very early papers it was already mentioned that the visual system seems to respond to temporal illuminance derivatives, rather than the luminance itself [50, 2], but the implications were not further developed.

## 4 Stabilized retinal images

When the retinal image is fully stabilized on the retina, perception fades in a few seconds. Any slight retinal slip generates edges perpendicular to the motion path [13, 39,37,38,2,20,21], see fig. 2.

**Fig. 2.**
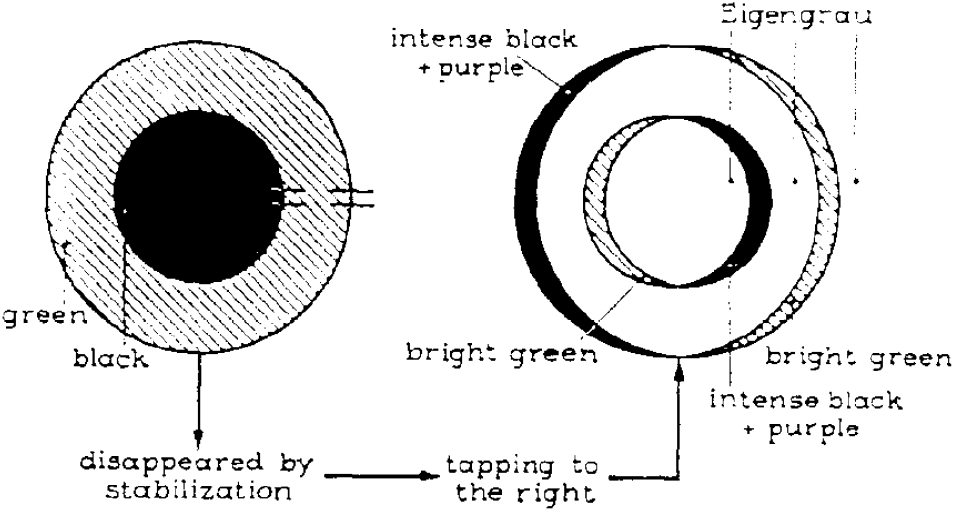
Stabilized retinal images: perception of ‘on’- and ‘off’-borders by slightly moving an object after the perceptlon has disappeared by stabilization. From [20].

It was an active research field in the early seventies. The classical explanation has been focused on adaptation, but the time difference model explains virtually all observed effects. Many prey animals move in a staccato manner (in combination with camouflage), in order to let the time difference fade their image away in the eye of the predator. Very familiar: ‘You see it when it starts to move’. The model also gives a new view on classical illusions. Even recenty the Troxler fading phenomenon, discovered in 1804, was explained as a result of neuronal adaptation [5]. The phenomenon is clearly well explained by the visual system taking the temporal derivative. The same holds true for the Anstit-Rogers ‘reverse phi motion’ illusion [1].

## 5 Directional micro-saccades induce frame difference edges on demand

Saccadic eye movements depend on attention driven tasks, but also on image content. Foulsham et al. found that when a stimulus image is rotated, also the saccadic orientation distribution follows the change in orientation, see figure 3. This happens in such a way that the saccades, in combination with the frame differencing, enhance edges perpendicular to the eye motion [18].

**Fig. 3.**
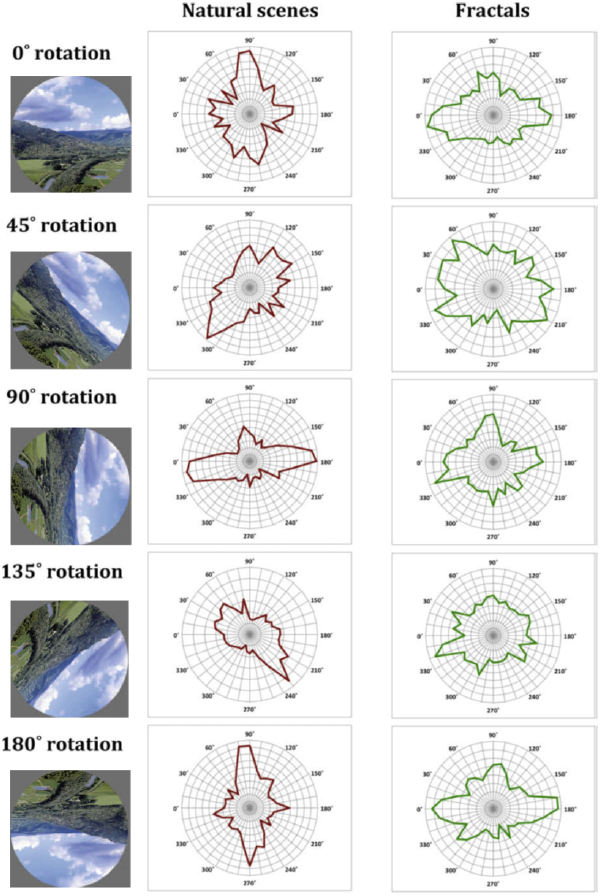
Saccadic eye movements depend on image content, and are preferentially made perpendicular to the major edges (figure adapted from [18]).

Foulsham et al. showed that binocular saccades exhibit a directional preference tendency along the epipolar line, supporting the enhancement of on-off difference induced edges perpendicular to the epipolar line for optimal depth estimation, while monocular saccades tyically are dependent on the local structure of edge orientations [18].

It was already noted by Gaarder in 1966 [19] that biases in micro-saccades directions depend on the characteristics of a given task, and give extra edges when contour recognition is weak or fails. The very short time course of micro-saccades (3-6 ms) gives a high 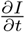 response, which is effective in generating edge information on higher cortical demand. Moreover, microsaccades show a clear preference for horizontal and vertical directions, the latter being less frequent in humans. Oblique directions are observed rather exceptionally [14, 32]. Research on microsaccades is rapidly increasing in the last decade (see an interesting citation analysis in [32]).

The new frame-difference model gives a new role for micro-saccades to be visually directed, as they are purposeful when the edge or texture information is incomplete or ambiguous. This challenges the classical notion that drift and micro-saccades are there to prevent fading [34].

## 6 Measurement of 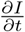 in real-time needs logarithmic time

Time differential structure requires multi-scale regularized derivative operators, just as in the spatial domain. They can be well represented as temporal Gaussian derivatives for pre-sampled stored temporal data. However, when we deal with real-time sampling, the time scale needs adjustment to be causal. A Gaussian temporal sampling cannot extent into the future. Koenderink showed from first principles that in a real-time sampling system the time axis needs to be reparametrized logarithmically [25, 49, 48]. The causal temporal Gaussian kernel and its first temporal derivative are given by

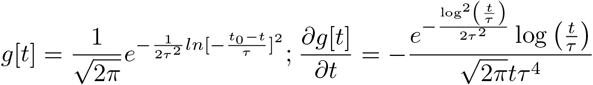

with *τ* the temporal scale. The last function is skewed, as can be seen in fig. 4b: DeValois carefully mapped the spatio-temporal receptive field sensitivity patterns of geniculate and cortical cells in the macaque monkey [10], and found indeed a skewed respons along the temporal axis of this real-time system (see fig. 4c).

**Fig. 4.**
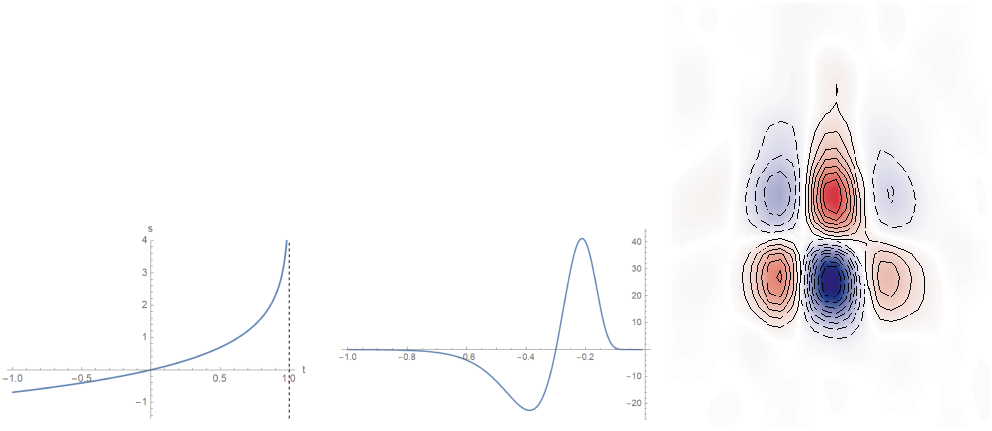
Causal time operator. Left: a. logarithmic reparametrization of time. Vertical axis: causal time, vertical axis: negative elapsed time (0 = the current moment). Middle: b. causal first order temporal derivative. Note the skewness. Right: c. spatio-temporal receptive field sensitivity profile of a V1 simple cell. Vertical axis: time, horizontal axis: space. Note the same skewness along the temporal axis (right figure from [10]).

## 7 Filling-in and internal representation

Taking temporal differences is a highly efficient energy saving and bandwidth limitation mechanism for vision. Only the information at locations of changed states needs to be transfered.

The price for the visual system to pay is a highly efficient and real-time filling-in process, active at all times. This phenomenon has been extensively studied and several models have been proposed, as reviewed by Komatsu [28], but the process is not yet fully understood. Recent fMRI studies trying to locate its activity have given conflicting results [44, 9]. In computer vision this process has been well developed as ‘image inpainting’, a fill-in restauration framework based on the pioneering work of [4], see fig. 5. A mathematical model, such as described in [4], might be a starting point to see if vision processes its filling-in algorithm in a similar way, given the powerful geometric computational capabilities of the visual cortex.

**Fig. 5.**
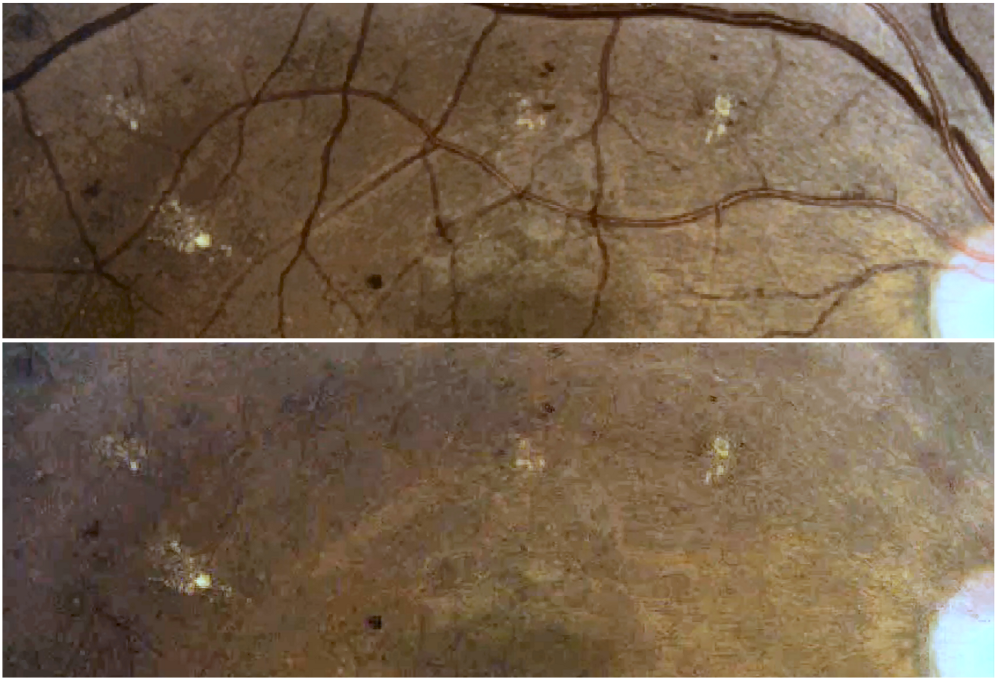
Removal of bloodvessels from a retinal image by image inpainting. Method: Mathematica 11.1 (Wolfram Inc.).

The filling-in process is closely related to our internal representation of the 3D outside world. It was convincingly shown by Nakayama that it is directly correlated with the perception of 3D depth [35]. A textured plane, occluded by a mask that uncovered only the corners, was seen through holes in the mask in two situations: before or behind the mask. The filling-in of the masked part of the plane is perceived markedly different in these situations: textured when in front, and void of texture when behind the mask.

## 8 3D-SLAM

At a larger scale, we do filling-in of our internal representation (a kind of painting of the complete scene, not inpainting of small patches) in our memory, which must be a truly 3D model of the world. The information through our optic nerve is limited. The number of RGCs, and thus the number of fibers in the optic nerve, is estimated to be a million, while the number of receptors is estimated to be around 150 million. The scanning is done with a looking-around foveated retina, which has a mosaic tiling with linear accuity decrease with excentricity. As an inspiration for internal ‘inpainting’ from the computer vision world, we discuss 3D Simultaneous Localization And Mapping (3D SLAM). With this technique a similar internal 3D representation is constructed by moving a sensor (camera or LIDAR) around in the scene, incrementally building up the complete framework, see fig. 6 and especially the corresponding movie. The code for such explorations runs in real-time on a CPU, and even on a modern smartphone [15] (on the other hand, a similar process must also run in the visual system of a navigating fruitfly [6]). This specialty field of robotics has benefitted greatly from Lie group theory, which is a key mathematical framework for geometrical transformations (and highly suitable for vision modeling).

**Fig. 6.**
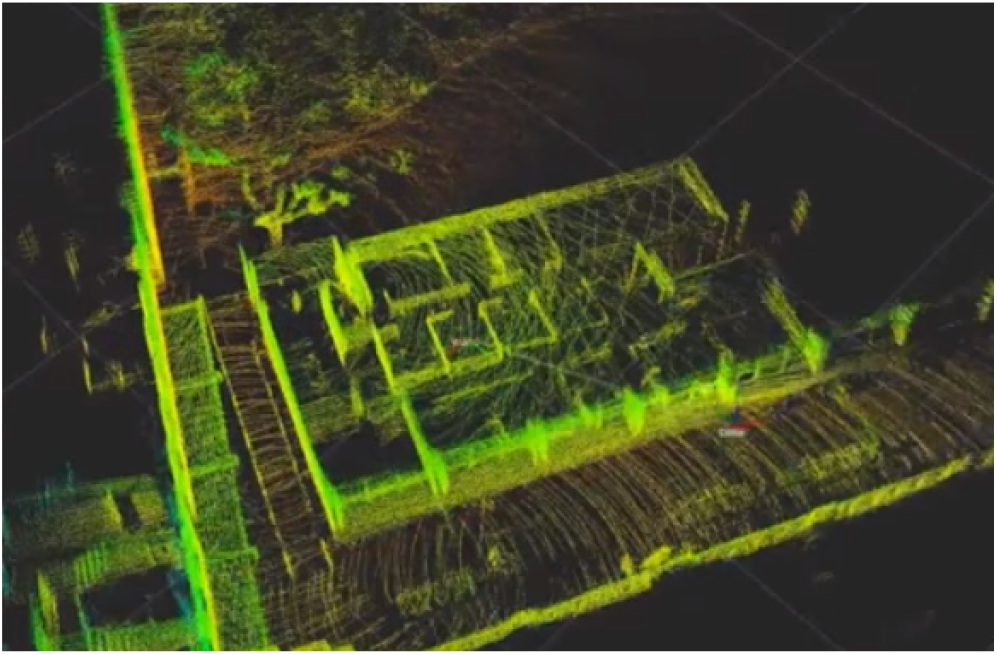
Wide indoor and outdoor real-time 3D SLAM (Simultaneous Localization and Mapping), exploiting a head-worn pulsed-laser LIDAR (Light Detection and Ranging) scanning device as analogon of a wondering foveated eye. From: Erik Nelson, UC Berkeley, Lawrence Berkeley National Laboratory, Berkeley. The effect is best appreciated by watching the movie: www.youtube.com/watch?v=08GTGfNneCI.

## 9 Discussion

The on-off pathway segregation is a fundamental design over virtually all visual systems, at the very basis of the processing pipeline. It is mediated by specialized bipolar cells, and found in the midget and parasol pathways, including its connecting stellate amacrine cells.

The possibility that the on- and off-channels might do a temporal subtraction, with one channel delayed, has been overlooked for a long time. Visual research has developed tremendously, but the time domain has been relatively underrepresented in vision measurements, which are predominantly based on morphology, electrophysiology, connectomics and biochemistry. The accessibility of recording sites for measuring relative cellular delay times of on-off pairs at cortical level is challenging.

Sometimes the terminology ‘on’ and ‘off’ is used to describe the positive and negative sensitivity areas in receptive field mappings of V1 simple cells, as in [30]. Here, however, the mapping is studied of cortical cells, which can be modeled by high order Gaussian derivative convolution kernels, with positive central sensitivity areas for even derivatives, and anti-symmetric kernels for odd derivatives.

The new insight gives a thorough explanation of the perceived phenomena with stabilized retinal images, and illusions like the Troxler fading and the Anstis phi motion.

It is quite counterintuitive to realize that we constantly look with a surveillance camera which is set to detect only differences. This requires an amazing continuous and real-time filling-in process. There are many indications we indeed have developed this process to great sophistication. The main driving evolutionary reason may be energy conservation, and the importance for survival of detecting changes (danger) by motions. That we are completely not aware of it, may be one of the reasons this finding may have been overlooked for so long.

## 10 Acknowledgements

This work is financed by the Hé Programme of Innovation Cooperation, which is (partly) financed by the Netherlands Organization for Scientific Research (NWO), dossier nr: 629.001.003. The project is also funded by the EU Marie Curie program ‘Metric Analysis For Emergent Technologies’ (MAnET), grant agreement no.: 607643. Video fig. 1 courtesy H.A.E.M. van Eldijk.

